# Priming lymphocyte responsiveness and differential T cell signaling in pediatric IBD patients with *Cannabis* use

**DOI:** 10.1101/2024.07.08.602495

**Authors:** Megan R. Sanctuary, Cinthia L. Hudacheck, Ashleigh J. Jones, Brittany V. Murphy, Nichole Welsh, Jost Klawitter, Edward J. Hoffenberg, Colm B. Collins

**Affiliations:** Conway Institute, School of Biomolecular & Biomedical Science, University College Dublin, Ireland; Department of Pediatrics, Division of Gastroenterology, Hepatology & Nutrition; Mucosal Inflammation Program; Digestive Health Institute, Children’s Hospital Colorado, University of Colorado, Anschutz Medical Campus, Aurora, CO; Department of Medicine, University of Colorado School of Medicine, Aurora, CO; Department of Anesthesiology, University of Colorado School of Medicine, Aurora, CO

**Keywords:** Cannabis, Inflammatory bowel disease, cytokine, pediatric

## Abstract

The prevalence of inflammatory bowel disease (IBD) has increased dramatically in recent years, particularly in pediatric populations. Successful remission with current therapies is limited and often transient, leading patients to seek alternative therapies for symptom relief, including the use of medical marijuana (*Cannabis sativa*). However, chronic cannabis use among IBD patients is associated with increased risk for surgical interventions. Therefore, determining the direct impact of cannabis use on immune modulation in IBD patients is of critical importance. Peripheral blood mononuclear cells of cannabis using and non-using pediatric IBD patients were phenotyped by flow cytometry and functionally assessed for their cytokine production profile. A phospho-kinase array was also performed to better understand changes in immune responses. Results were then compared with serum phytocannabinoid profiles of each patient to identify cannabinoid-correlated changes in immune responses.Results demonstrated elevated levels of a myriad of pro-inflammatory cytokines in users versus non-users. Differences in signaling cascades of activated T cells between users and non-users were also observed. A number of anti-inflammatory cytokines were inversely correlated with serum phytocannabinoids. These results suggest that cannabis exposure, which can desensitize cannabinoid receptors, may prime pro-inflammatory pathways in pediatric IBD patients.

**Article Summary:** This observational study examines the impact of chronic cannabis use on peripheral immune cell function in adolescent IBD patients from Children’s Hospital Colorado. Cannabis users displayed altered T cell phenotype, increased pro-inflammatory cytokine release and dephosphorylation of protective protein kinases.

## Introduction

Thirty percent of adolescent and young adult IBD patients at Children’s Hospital Colorado report having used marijuana to treat their disease symptoms despite the absence of credible evidence for therapeutic benefit ^1^. More broadly, cannabis has gained interest among inflammatory bowel disease patients due to its capacity to suppress pain, reduced gut motility, improve appetite and stimulate weight gain ^2^. This has been borne out in a number of small short-duration clinical studies ^3^. While these studies have reported improvement in subjective clinical indicators, little evidence of improvement by empirical markers of intestinal disease or mucosal healing exists. Moreover, concerns exist regarding the long-term impact of cannabis use, as it is associated with a five-fold increase in surgical risk among Crohn’s disease (CD) patients ^4^.

This concern is further amplified in adolescent IBD patients, where any symptomatic relief provided by cannabis use must outweigh well-established negative implications of cannabis on cognitive function within the still developing brain ^5^. Cannabis exposure during adolescence can lead to neurocognitive disadvantages, including in relation to attention and memory, that can persist beyond abstinence ^6^.

Within the intestine, both endogenous (endo-) or plant-derived (phyto-) cannabinoids signal via two G-protein coupled receptors, CB_1_R and CB_2_R (encoded by *CNR1* and *CNR2*) and by non-canonical receptors such as GPR55, PPARα and TRPV1 ^2^. CB_1_R are largely localized to extrinsic neurons and vagal efferents as well as epithelial cells where they regulate intestinal permeability ^7^. In contrast, CB_2_R is mainly expressed on immune cells, specifically neutrophils, macrophages, T and B cells. This system is intended to respond to endocannabinoids, including anandamide (AEA) and 2-arachidonoylglycerol (2-AG), that are synthesized on demand from membrane precursors. However, these receptors can also be triggered by the over 100 active ingredients present in the cannabis plant including the weak agonist Δ9-tetrahydrocannabinol (THC), the inverse agonist cannabidiol (CBD) and a range of less highly expressed agonists, inverse agonists and allosteric modulators. It is because of the latter that whole plant extracts are thought to have greater efficacy due to a synergistic or “entourage” effect though there is limited evidence that this is the case.

Expression of *CNR1* and *CNR2* mRNA are downregulated in the ilea of CD patients ^8^ even in the absence of exogenous phytocannabinoid signaling and so it remains to be seen what impact chronic receptor activation might have on an already disrupted endocannabinoid tone. Furthermore, whereas a number of acute murine colitis studies have reported protective effects of CB_2_R activation ^9^, CB_2_R blockade attenuated inflammation in a chronic murine ileitis model of human CD ^8^ suggesting that cannabinoid receptor activation elicits differing responses depending on acute or chronic inflammatory conditions.

A number of cannabis-based medications have already been approved for clinical use including dronabinol, a synthetic THC, for the treatment of anorexia associated with weight loss in AIDS patients ^10^. Nabilone, an orally active synthetic cannabinoid is FDA-approved for the treatment of chemotherapy-induced nausea and vomiting ^11^ and CBD is approved for the treatment of seizures associated with Lennox-Gastaut syndrome or Dravet syndrome in patients 2 years of age and older ^12^ paving the way for potential approval for IBD.

In summary, while cannabis use is prevalent among IBD patients, physicians have insufficient evidence to recommend cannabis as a form of complementary care. It is essential that we better understand at a basic cellular level, the impact of chronic cannabis use on immune function in patients with chronic immunological disorders as distinct from acute cannabinoid exposure or during acute inflammation. In order to begin to understand the impact of cannabis use on the immune function of adolescent IBD patients, we collected peripheral blood from patients, both using and non-users, and subject them to both phenotypic and functional analysis. We performed HPCL-based quantification of serum phytocannabinoids in order to correlate immunological changes with phytocannabinoid expression. Based on our initial findings, we then subjected isolated leukocytes to quantification of changes in phosphorylation of key signaling molecules in order to identify potential mechanisms that may account for the differences in immune responses seen between leukocytes obtained from users and non-users.

## Methods

Adolescent (13 to 23 years old) IBD patients at Children’s Hospital Colorado were enrolled as part of a larger prospective study evaluating cannabis exposure in the adolescent population ^1^. Patients were recruited, independent of cannabis use, by a research coordinator during either outpatient visit, inpatient admission or medication infusioSSn visit (**Figure 1**).

**Figure 1.**
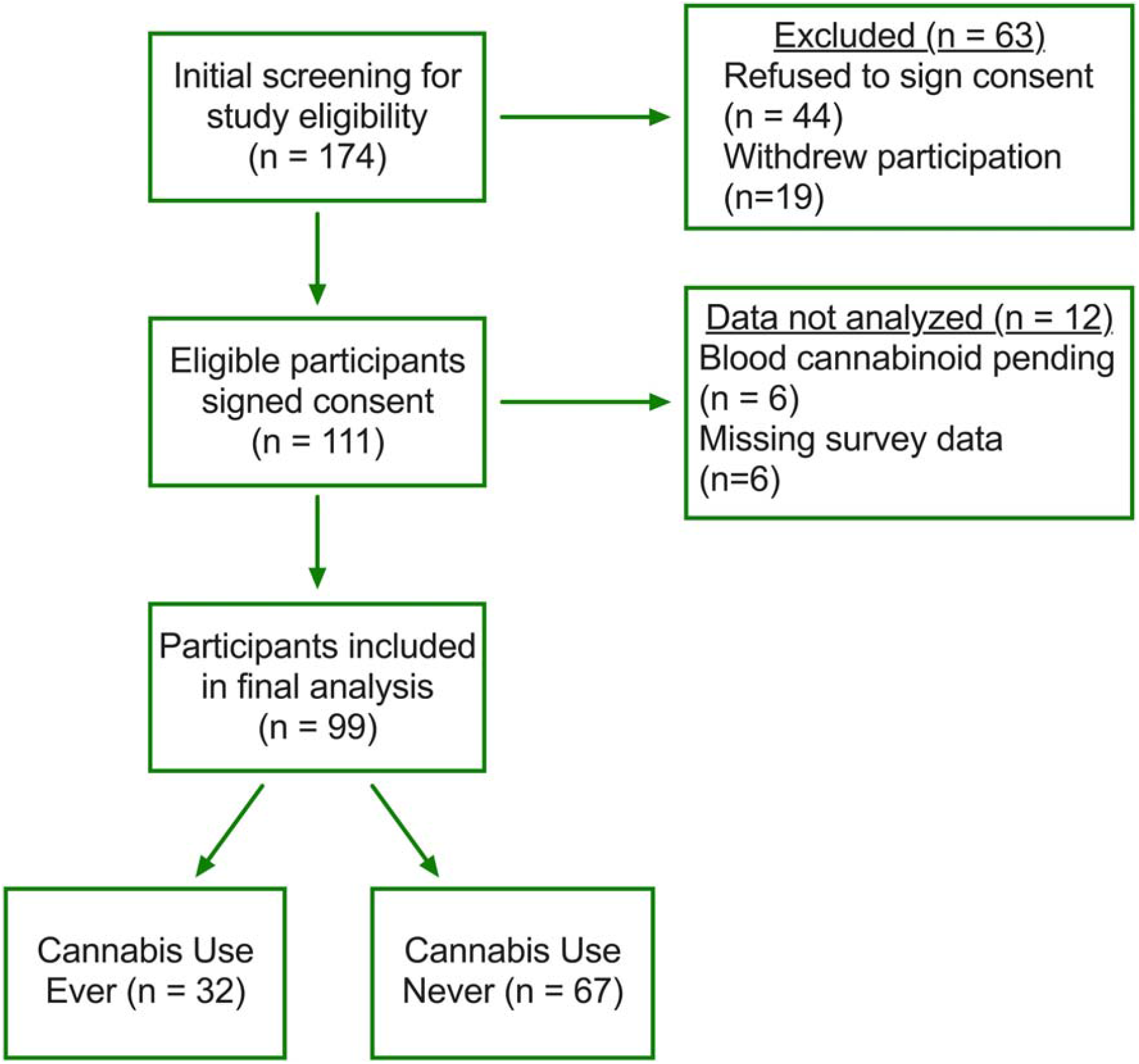
Study design schematic. Flow chart outlining study design and participant recruitment, enrollment, and analysis. Of the 174 IBD patients screened for enrollment, 64% were consented for participation. Of the 111 patients who completed the study, 89% were included in the final analysis. Based on the final analysis, 67% never used cannabis while 32% of patients had ever used cannabis.

### Sample Collection

Deidentified whole blood samples were collected from pediatric IBD patients in heparinized glass CPT tubes and centrifuged according to manufacturer’s instructions. Separately, 1ml of serum was collected in a glass container for subsequent cannabinoid quantification provided by iC42 as described ^13^ (Aurora, CO). The peripheral blood mononuclear cell (PBMC) fraction was resuspended in 20% DMSO and 80% inactivated Fetal Calf Serum, adjusted to 1×10^7^ cells/ml, aliquoted and frozen in standard cryovials. After 24hrs of slow freezing at -80°C, samples were transferred to liquid nitrogen for long-term storage.

### Flow Cytometry

Samples were thawed and rested for 1hr in RPMI with 10% FBS at 37°C prior to restimulation in PMA (50ng/mL), Ionomycin (1μg/mL) and brefeldin A (10μg/mL) for 4hr at 37°C. Cells were then fixed, permeabilized and stained according to the eBioscience (San Diego, CA) FoxP3/Transcription Factor Staining buffer set instructions. Samples were stained using fluorescently conjugated antibodies for CD3 (OKT3), CD4 (RPA-T4), CD25 (BC96), CD127 (A019D5), FoxP3 (206D), IFNγ (B27), IL-10 (JES3-9D7) and IL-17A (BL168; Biolegend San Diego, CA) on a BD LSRII and analyzed on Flowjo software using appropriate negative and fluorescent-minus one controls.

### Cytokine Secretion

Thawed PBMC were counted and plated in triplicate on 48-well U-bottomed assay plates pre-coated with anti-CD3 (OKT3; 10μg/mL) at 1×10^6^ cells/well. Cells were incubated for 24hr at 37°C in media containing anti-CD28 (CD28.2; 500ng/mL). Samples were transferred to microcentrifuge tubes, centrifuged at 1000g for 5 minutes and supernatants transferred to fresh cryovials for subsequent storage prior to analysis. Supernatant cytokines were assayed using the Bio-Plex Human Cytokine 17-plex assay system (Biorad, Hercules CA) according to manufacturer’s instructions.

### Phospho-kinase Profiling

Using the Human Phospho-kinase Antibody Array™ (R&D Systems), we assayed site-specific changes in phosphorylation of 43 human kinases. PBMC were thawed and rested for 4hr at 37°C prior to stimulation with anti-CD3/CD28 (10μg/mL) for 5 minutes. Cells were then placed on ice with ice-cold PBS added prior to lysis and blotting according to manufacturer’s instructions.

### Statistics

Statistical analyses were performed using the ROUT test to identify outliers followed by the nonparametric Mann-Whitney U test using Prism 8 software (Graphphad, La Jolla, CA). Data were expressed as mean ± standard error of the mean (SEM). Correlation analysis was performed using Pearson correlation coefficients. Statistical significance was set at *P*<0.05.

### Ethical Considerations

Study protocols were approved by the Ethical Committees of Children’s Hospital Colorado. All patients and controls gave their written and informed consent to participate according to Colorado Multiple Institute Research Board protocol #15-0416.

## Results

### Patient recruitment

We identified 32 patients (**Figure 1**) who reported having ever used cannabis, with 21 patients testing positive for serum phytocannabinoids at the time of sample collection. The most consistent phytocannabinoid measured was the THC metabolite 11-nor-9-carboxytetrahydrocannabinol (THC-COOH), with a mean serum concentration of 36.6 ± 10.4 ng/ml across 20 patients followed by the metabolite THC-glucuronide (114 ± 35.8 ng/ml **Supplemental Figure 1**) present in 19 patients. There were no significant differences in terms of age, gender, race, disease type or physician global assessment (**Table 1**). Among cannabis users, there was an even split between those who used cannabis less than weekly or more than weekly but within those who used weekly or more, more than twenty percent used twice a day or more (**Table 2**).

**Table 1.**

|  | All Participants<br>(N = 99) | Never Used Cannabis<br>(N = 67) | Ever Used Cannabis<br>(N = 32) | Statistic:<br>P-value |
| --- | --- | --- | --- | --- |
| Years of Age<br>(mean±SEM) | 16.2 ± 2.2 | 15.9 ± 2.2 | 17.0 ± 1.9 | t (97) = 2.6;<br>P=0.011 |
| Male (N, %) | 56, 57% | 37, 55% | 19, 59% | $\chi^2(1)=0.015$;<br>P=0.70 |
| Race (N, %)<br>• White/Caucasian<br>• Other | 80, 82%<br>18, 18% | 56, 84%<br>11, 16% | 24, 77%<br>7, 23% | $\chi^2(1)=0.54$;<br>P=0.46 |
| Disease Type<br>• UC<br>• CD<br>• Unknown | 27, 27%<br>62, 63%<br>10, 10% | 21, 31%<br>40, 60%<br>6, 9% | 6, 19%<br>22, 69%<br>4, 13% | $\chi^2(1)=1.81$;<br>P=0.404 |
| Physician Global Assessment (N, %)<br>• Inactive/mild<br>• Moderate/severe | 82, 83%<br>17, 17% | 55, 82%<br>12, 18% | 27, 84%<br>5, 16% | $\chi^2(1)=0.08$;<br>P=0.78 |
| Detected Serum Cannabinoids (N, %) | 16, 16% | 0, 0% | 16, 50% | $\chi^2(1)=40.0$;<br>P<0.001 |

**Table 2.**

|  | Frequency of Use | N (%) |
| --- | --- | --- |
| Use less than weekly | • No use<br>• < Once per month<br>• Once per month<br>• ≥ Twice per month<br>○ Total | 2 (7%)<br>9 (31%)<br>2 (7%)<br>1 (3%)<br>14 (48%) |
| Use weekly or more | • < Once per week<br>• ≥ Twice per week<br>• Once per day<br>• ≥ Twice per day<br>○ Total | 3 (10%)<br>3 (10%)<br>3 (10%)<br>6 (21%)<br>15 (51%) |

### Increased cytokine secretion by lymphocytes associated with cannabis use

We evaluated the frequency of CD4^+^ T cell subsets from the peripheral blood of adolescent IBD patients based on expression of cytokines interferon gamma (IFN?), interleukin-10 (IL-10) and interleukin 17 (IL-17) using the gating strategy outlined in **Figure 2A**. There were no significant differences in the frequency of total CD4^+^ T cells between non-users and cannabis users (21.1 ± 2.3 in non-users to 21.7 ± 0.9 in users; not shown). However, cannabis use was associated with an increased frequency of IFN?^+^ cells (from 6.5 ± 0.8 in non-users to 11.6 ± 1.7 in users; P<0.01) expressed as percent of CD4^+^ cells. The frequency of IL-10^+^ cells increased (from 1.1 ± 0.2 to 2.8 ± 0.9; P<0.05). Similarly, the frequency of IL-17^+^ Th17 cells increased significantly from non-users to users (2.4 ± 0.4 to 5.13 ± 1.27; P<0.05). Representative zebra plots illustrate the effect of cannabis use on the production of cytokines between users and non-users with fluorescence minus one controls included for the purpose of gating. Therefore, while we did observe a significant increase in the frequency of Th1 and Th17 cells in PBMC of cannabis users, we also observed an increase in IL-10 producing T cells suggesting that lymphocytes from cannabis users were more primed overall.

**Figure 2.**
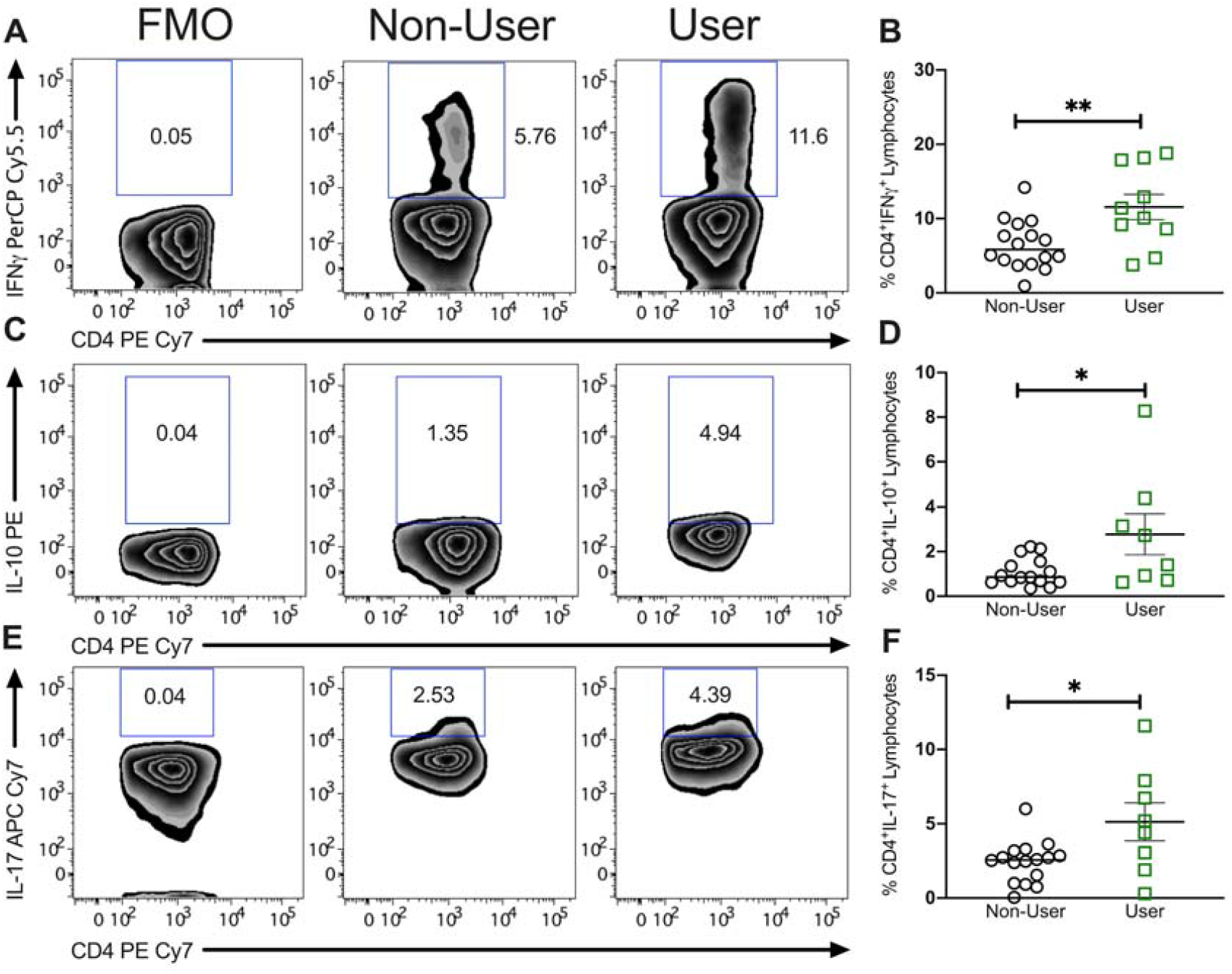
Proportion of circulating pro-inflammatory Th17 cells is higher in pediatric IBD patients using cannabis compared to non-users. Representative zebra plots of intracellular cytokine staining for (A) IFNγ, (C) IL-10 and (E) IL-17 including fluorescence minus one (FMO) gating control. Scatter plots of percentage of cytokine positive CD4^+^ T cells from PBMC of non-user and cannabis using adolescent IBD patients showing the significant increase in (B) IFNγ, (D) IL-10 and (F) IL-17 by patients within the ‘user’ group. Results represent mean±SEM for N ≥ 10 individual patients. *P<0.05, **P<0.01.

### Cannabis users displayed an increased frequency of FoxP3^+^ cells in circulation

In order to analyze expression of regulatory T cell subsets, CD4^+^ T cells were initially subdivided based on expression of CD25 (IL-2R) and CD127 (IL-7R) into total CD25^High^CD127^Low^ Tregs and non-Tregs CD25^Low^CD127^+^ and CD25^Neg^CD127^Neg^ (**Figure 3A)**. This failed to identify a significant difference in frequency of these cell populations. Interestingly, when we instead analyzed expression of the canonical Treg transcription factor FoxP3, we identified a significant increase in expression of CD4^+^ FoxP3^+^ cells among cannabis users relative to non-users (4.3 ± 0.5 to 7.16 ± 0.8; P<0.01; **Figure 3D**). Expression of FoxP3 defines Tregs with high regulatory activity, and as demonstrated in **Figure 3E**, FoxP3 expression is not restricted to CD25^High^CD127^Low^ Tregs. Thus, while expression of CD25^High^CD127^Low^ Tregs including high and lowly suppressive Treg subpopulations was unchanged in cannabis users, expression of FoxP3^+^ cells were increased in peripheral blood.

**Figure 3.**
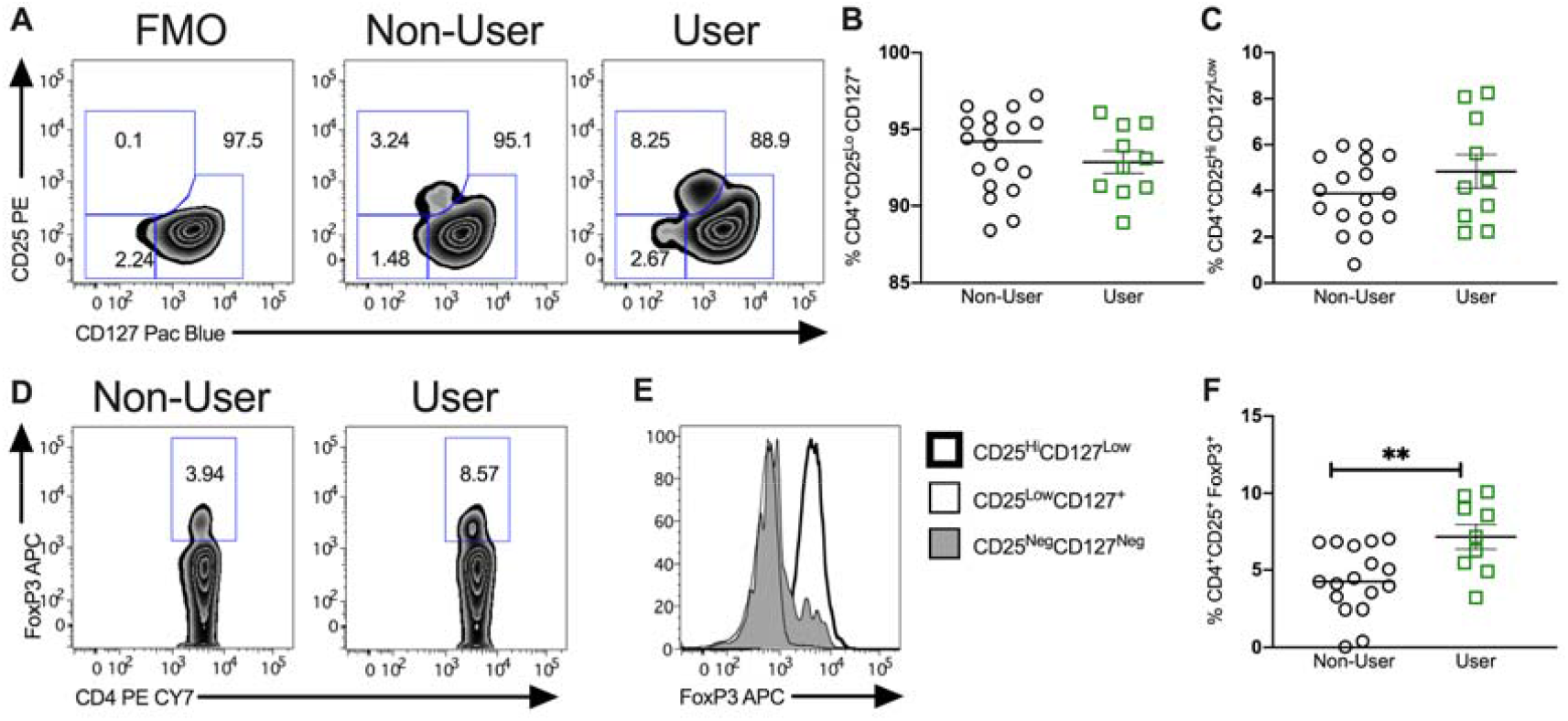
Impact of cannabis use on circulating regulatory T cells. (A) Representative zebra plots of flow cytometric characterization of the CD4^+^ T cell fraction of previously frozen peripheral blood mononuclear cells. (B) Relative frequency of CD4^+^CD25^lo^CD127^+^ T cells between non-users and users. (C) Relative frequency of CD4^+^CD25^Hi^CD127^Low^ T cells between non-users and users. (D) Representative zebra plots of FoxP3^+^ CD4^+^ T cells from adolescent IBD patients using or not using cannabis with representative histogram for FoxP3 (E) and quantification of regulatory T cell frequency (F). Results represent mean±SEM for N ≥ 10 patients, **P<0.01.

### T cell-associated cytokine secretion is unaffected in cannabis users

Having identified differences in the frequency of T cell subsets, we next assessed the cytokine secreting function of those cells. Isolated PBMC were subjected to 24hr T cell-specific activation and secreted cytokines were assessed by multiplex cytometric bead array. Despite significant differences observed in the frequency of T cell subsets, no significant differences were seen in the overall levels of Th1 (IL-12), Th2 (IL-4), (B) Th17 (IL-17) or Treg (IL-10) associated cytokines between IBD patients who used cannabis and those who did not (**Supplemental Figure 2**). Thus, while cannabis use is associated with an alteration in the relative abundance of CD4^+^ T cells within the PBMC fraction, this does not appear to be adequate to elicit changes in T cell-associated cytokine secretion.

### Enhanced bystander responses of leukocytes obtained from cannabis users

We next broadened our studies to investigate the impact on additional pro and anti-inflammatory cytokines. In cultured supernatants, there was a significant increase in the secretion of IL-1β (**Figure 4A**) among cannabis users (77.24 ± 26.18 vs 486.7 ± 172.1 pg/ml; P<0.05), differences in IL-6 though considerable failed to reach statistical significance (148.7 ± 37.67 vs 505.0 ± 185.2 pg/ml; P=0.1) while MCP-1 was also significantly elevated in stimulated PBMC from cannabis using adolescents relative to non-users (226.1 ± 66.16 vs 916.0 ± 222.1 pg/ml; P<0.005). In a similar way, secretion of TNFL (**Figure 4B**; 8.71 ± 1.4 vs 18.1 ± 4.1 ng/ml; P<0.05), IL-8 (23.5 ± 7.1 vs 90.6 ± 16.9 ng/ml; P<0.005) and MIP-1β (2.5 ± 4.5 vs 3.7 ± 1.0 ng/ml; P=0.2) were all elevated though MIP-1β failed to reach statistical significance. Likewise secretion of IL-7, G-CSF (granulocyte colony-stimulating factor) and GM-CSF (granulocyte-macrophage colony-stimulating factor) all failed to be significantly altered (**Figure 4C**).

**Figure 4.**
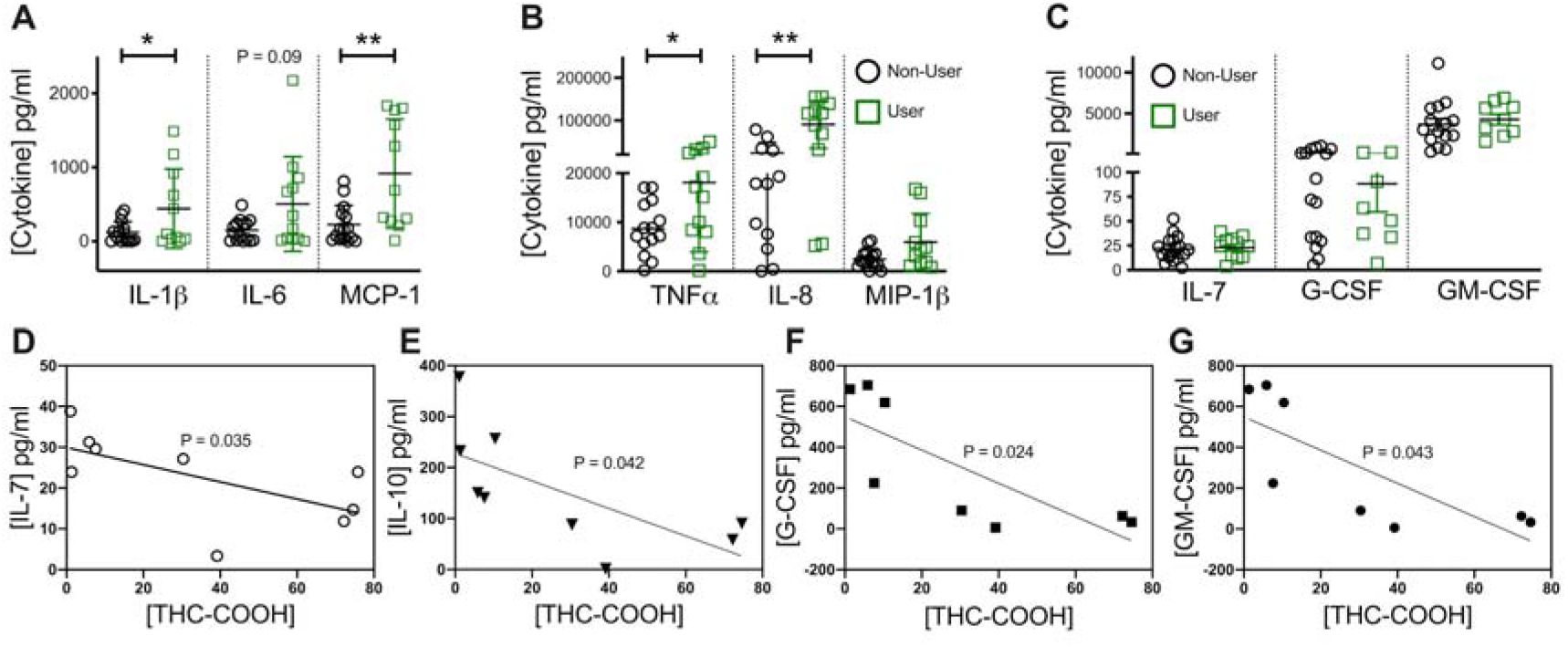
Production of inflammatory cytokines is elevated in stimulated PBMCs from pediatric IBD patients using cannabis compared to non-users. Multiplex analysis of cytokine production by previously frozen PBMC identified a significant increase in non-T cell-associated pro-inflammatory cytokines (A) IL-1β, IL-6 and monocyte chemoattractant protein-1 (MCP-1) as well as (B) TNFα and IL-8 but not in (C) IL-7, G-CSF and GM-CSF after 24h restimulation with anti-CD3/CD28 antibodies. Correlation analysis with phytocannabinoid THC-COOH identified a significant inverse correlation between cannabinoid levels and (D) IL-7, (E) IL-10, (F) G-CSF and (G) GM-CSF. Results represent mean±SEM for N ≥ 8. *P<0.05, **P<0.01.

To better understand the variability in cytokine responses, we performed correlation analysis between individual patient cytokine responses and their serum phytocannabinoids. Based on Pearson correlation analysis, IL-10 expression inversely correlated with serum levels of THC (R = -0.82; P<0.05) and its metabolite THC-COOH (R = -0.71; P<0.05; **Figure 4D**). Similarly IL-7 levels inversely correlated with THC-COOH (R = -0.79; P<0.01; **Figure 4E**) as did G-CSF (R -0.80 P<0.05; **Figure 4F**) and GM-CSF (−0.65 P<0.05; **Figure 4G**). As such, this data identifies a significant increase in pro-inflammatory cytokine production upon T cell reactivation in cannabis users along with an inverse correlation between circulating THC levels and anti-inflammatory IL-10.

### Altered signaling response in PBMC from cannabis users

To evaluate potential mechanisms driving the alterations in cytokine responses from PBMC of cannabis users relative to non-users, PBMC were subject to brief T cell activation and subsequent phosphorylation of key signaling proteins detected by phospho-array. This identified seven differentially phosphorylated phospho-kinases, six of which expressed a decrease in phosphorylation in cannabis users relative to non-users including Hck from 1 ± 0.05 to 0.8 ± 0.03 (P<0.05; **Figure 5**), Fyn from 1 ± 0.1 to 0.7 ± 0.06 (P<0.01), FAK from 1 ± 0.09 to 0.7 ± 0.15 (P<0.05), ERK1/2 from 1 ± 0.05 to 0.7 ± 0.04 (P<0.001), Stat5a from 1 ± 0.09 in non-users to 0.6 ± 0.09 fold in users (P<0.005), β-catenin from 1 ± 0.09 to 0.7 ± 0.27 (P<0.05) and p53 at residue S15 from 1 ± 0.08 to 0.7 ± 0.05 (P<0.05). In contrast phosphorylation of HSP27 at position Ser78/82 was increased in cannabis users (1 ± 0.07 to 1.5 ± 0.22; P<0.05) indicative of an increase in tolerance to noxious stimuli. Taken together, we observed alterations in a number of key phospho-kinases in cannabis user-derived PBMC relative to non-users which may have implications for cell survival, trafficking and proliferation.

**Figure 5.**
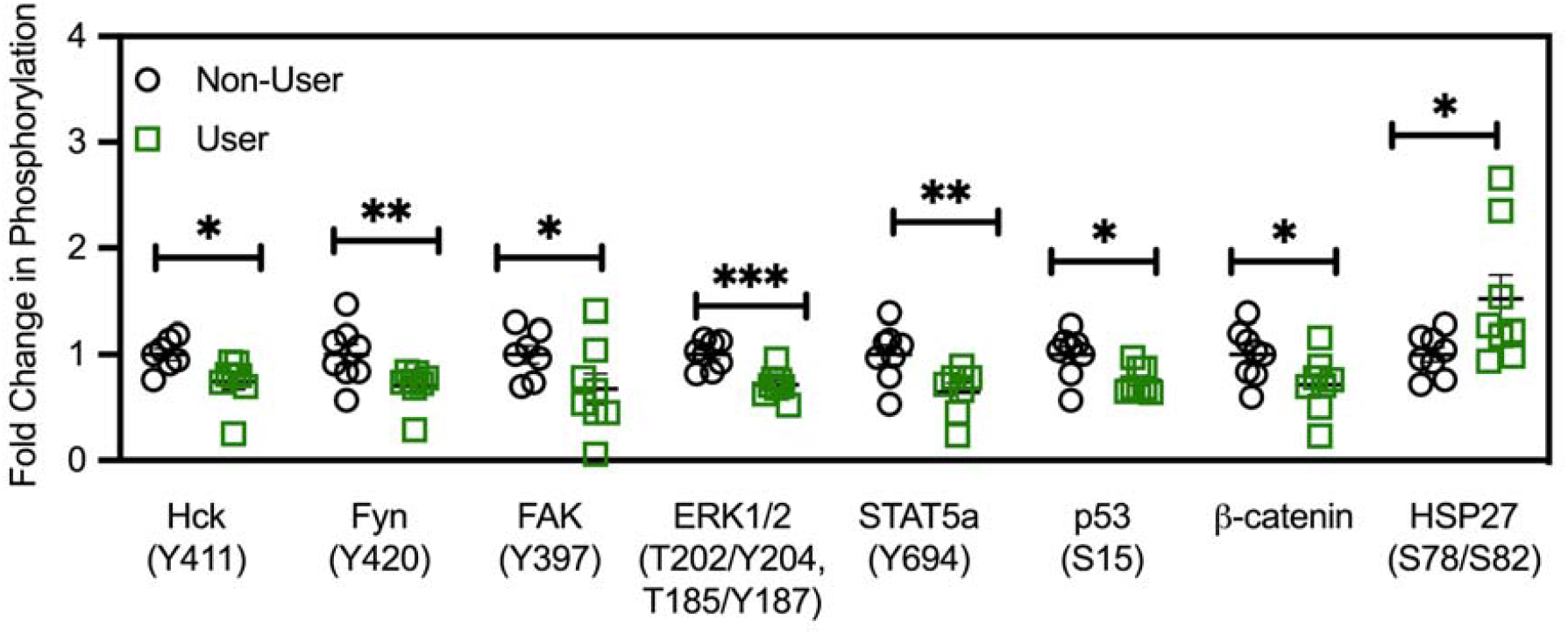
Cannabis-associated changes in proteome profile. Comparison of proteome profile of TCR-activated PBMC from adolescent IBD patients demonstrate a significant decrease in phosphorylation at known activation sites for Hck, Fyn, FAK, ERK1/2, STAT5a, p53 and β-catenin with a significant increase in activation-associated phosphorylation seen at HSP27 only. Results represent mean±SEM for N = 8 age and gender-matched patients per group. *P<0.05, **P<0.01, ***P<0.001.

## Discussion

Previous studies have demonstrated that *in vitro* cannabinoid administration can elicit anti-inflammatory effects on colonic explants from IBD patients and cultured intestinal epithelial cells ^7^ supporting the need for further investigation. This study builds on those findings, by assessing the impact of *in vivo* cannabis use on IBD patient immune responses among an adolescent population.

Using flow cytometry to characterize T cell subsets, we first identified similar T cell frequencies in peripheral blood of pediatric IBD patients compared with those previously described ^14^. Interestingly cannabis use was associated with an increased frequency of IFN?^+^ and IL-17^+^ T cells compared with non-users. Previous studies in adult ulcerative colitis (UC) patients identified an association between an increase in circulating Th1 and Th17 cells with more active disease ^15^. While we failed to observe significant clinical differences in disease severity between users and non-users, this early divergence drive worsening outcomes in later stages of the disease.

In contrast, we also observed an increase in FoxP3^+^ CD4^+^ T cells in the peripheral blood, consistent with previous studies using THC treatment in a rhesus macaque model of HIV ^16^. IBD patients exhibit an increase in peripheral blood Tregs frequency during disease remission ^17^ associated with an decreased Treg expression at mucosal sites ^17^. It is plausible therefore that the relative increase seen in cannabis users may reflect less active disease.

While it is difficult to draw conclusions regarding the overall impact of these changes in the T cell compartment, nevertheless it appears that for better or worse, cannabis use alters immune phenotype in adolescent IBD patients.

We next assessed functional differences between PBMC from users and non-users. TCR activation triggered a knock-on effect promoting greater pro-inflammatory cytokine secretion from PBMC of cannabis users. TCR activation induces cytokine production including IL-1β, TNFL and IL-8 indirectly from monocytes present in the PBMC via close contact ^18^. Once again, this is concerning given that increased immune reactivity is associated with more rapid disease progression and increased hazard ratio ^19^.

A clear limitation of this observational study is the variable cannabis exposure experienced by patients in the “user” group. Much of the varsiation might also be obtained in an interventional study also, given the highly variable nature of cannabis pharmacokinetics ^20^. Nonetheless, to determine if the observed effects on cytokine production could be linked to serum cannabinoid levels, we performed correlation analysis of cytokines with serum phytocannabinoids. Upon ingestion, THC is metabolised to 11-hydroxytetrahydrocannabinol (11-OH-THC) and THC-COOH with the latter found at significantly higher concentrations in plasma 24hrs post ingestion making it a useful surrogate marker for THC exposure ^21^.

We observed an inverse correlation between THC and IL-10 (and its active metabolite THC-COOH), while IL-7, G-CSF and GM-CSF were inversely correlated with the more stable THC-COOH only. IL-10 deficiency is used as a murine IBD model of spontaneous colitis and clinical trials have demonstrated efficacy of recombinant IL-10 administration in mild to moderate Crohn’s ^22^. While clinical use of IL-10 may be limited due to the highly localized nature of its action, IL-10 is clearly an important anti-inflammatory cytokine and so it is concerning that the production of IL-10 is inversely correlated to patients serum phytocannabinoid levels. Similarly, IL-7 has a role in epithelial integrity and T cell memory stability during IBD ^23^. IL-7 levels are increased three-fold in serum of pediatric IBD patients relative to age-matched controls ^24^. Mucosal IL-7 decreased with worsening mayo scores in UC patients, albeit likely due to a loss of goblet cells ^25^. Similarly, levels of G-CSF, a pleiotropic immunomodulatory cytokine which is already expressed at a lower level in IBD patient intestinal mononuclear cells ^26^ inversely correlated with THC-COOH. Pre-treatment with G-CSF suppresses immune activation of PBMC ^26^ and long-term supplementation of G-CSF induced remission in a pediatric patient with UC-like IBD linked to glycogen storage disease Ib ^27^. Reduced GM-CSF may also have a negative consequence for pediatric IBD patients given the therapeutic efficacy of Sargramostim, the recombinant GM-CSF ^28^. Loss of the ligand would represent a further blow to this immunoregulatory system which sees expression of the GM-CSF receptor (CD116) significantly decreased in IBD patients ^29^.

In order to delve deeper into the mechanisms driving the increased immune response of PBMC from cannabis users, we profiled changes in cell signaling using a phospho-kinase proteomic profiler array. We identified decreased expression of phosphorylated β-catenin and downstream phosphorylated PLC-L1 in cannabis users compared with non-users. β-catenin serves as a negative regulator of T cell activation through selective interference of LAT-PLC-L1 phosphorylation ^30^ which may have contributed to the increased cytokine responses. Previous studies in colorectal cancer where CB_2_R is a poor prognostic factor, have demonstrated suppression of β-catenin function in response to CB_2_R activation in colon cancer cell lines ^31^ further corroborating our findings. We hypothesize that loss of this negative regulation enhances T cell activation driving increased cytokine secretion by monocytes. Heightened sensitivity of the TCR may be insufficient alone to trigger a disease flare in patients but would likely exacerbate symptoms upon relapse.

Cannabinoids activate p53 during neuronal cell death ^32^ but within peripheral blood lymphocytes, CB_2_R activation downregulates p53 ^33^ similar to our findings. In addition, p53 may be impacted indirectly, as a consequence of a dampened upstream pathway such as Fms-like tyrosine kinase (Flt3) signaling, which is also associated with inactivation of p53 ^34^. Also dephosphorylated in our study is hematopoietic cell kinase (Hck), an src family member activated upstream of ERK and STAT5 phosphorylation ^35^. It is preferentially expressed by myeloid cells ^36^ and can also be activated by Flt3 ^37^ which we have previously demonstrated to be an important anti-inflammatory pathway in a pre-clinical IBD model. TCR stimulation increases ERK1/2 phosphorylation, which in turn drives progression of CD4^+^ T cells lineage determination. The relative decrease in ERK phosphorylation seen in cannabis users in our study is consistent with previous work which demonstrated that prior activation of CB_2_R can suppress subsequent induction of ERK1/2 phosphorylation in whole blood cells ^38^.

TCR activation also induces tyrosine phosphorylation of focal adhesion kinase (FAK) at position 397. FAK is sensitive to Src family kinase inhibition ^39^ suggesting that the decreased phosphorylation seen in activated PBMC from cannabis users reflects a downregulation of a common signaling pathway with ERK, Fyn and Hck. In addition to activating downstream ERK, Fyn contributes to proximal TCR signaling with loss of Fyn leading to primary T cells becoming activated more rapidly and to produce more cytokines ^40^. The increased cytokine responses seen in our studies may therefore reflect in part a decreased phosphorylation of Fyn at its activation side at tyrosine 420 associated with chronic cannabis use. Though an effect of cannabis on 420 has not been described for human immune cells previously.

## Conclusion

Our findings suggest that cannabis use among adolescent IBD patients alters T cell signaling to promote pro-inflammatory cytokine release from PBMC. This primed immune system may exacerbate disease flares and drive the increased surgical intervention seen in adult IBD patients who use cannabis ^4^.

## Acknowledgement of Funding

This work was funded by the Crohn’s & Colitis Foundation SRA (CBC), the UCD Ad Astra Program (CBC), the Colorado Department of Public Health and Environment (CDPHE; CBC/EJH), and Colorado Clinical Translational Science Institute (CBC) which is supported by NIH/NCATS Colorado CTSA Grant Number UL1 TR002535.

## Supplemental Figures

**Figure S1.**
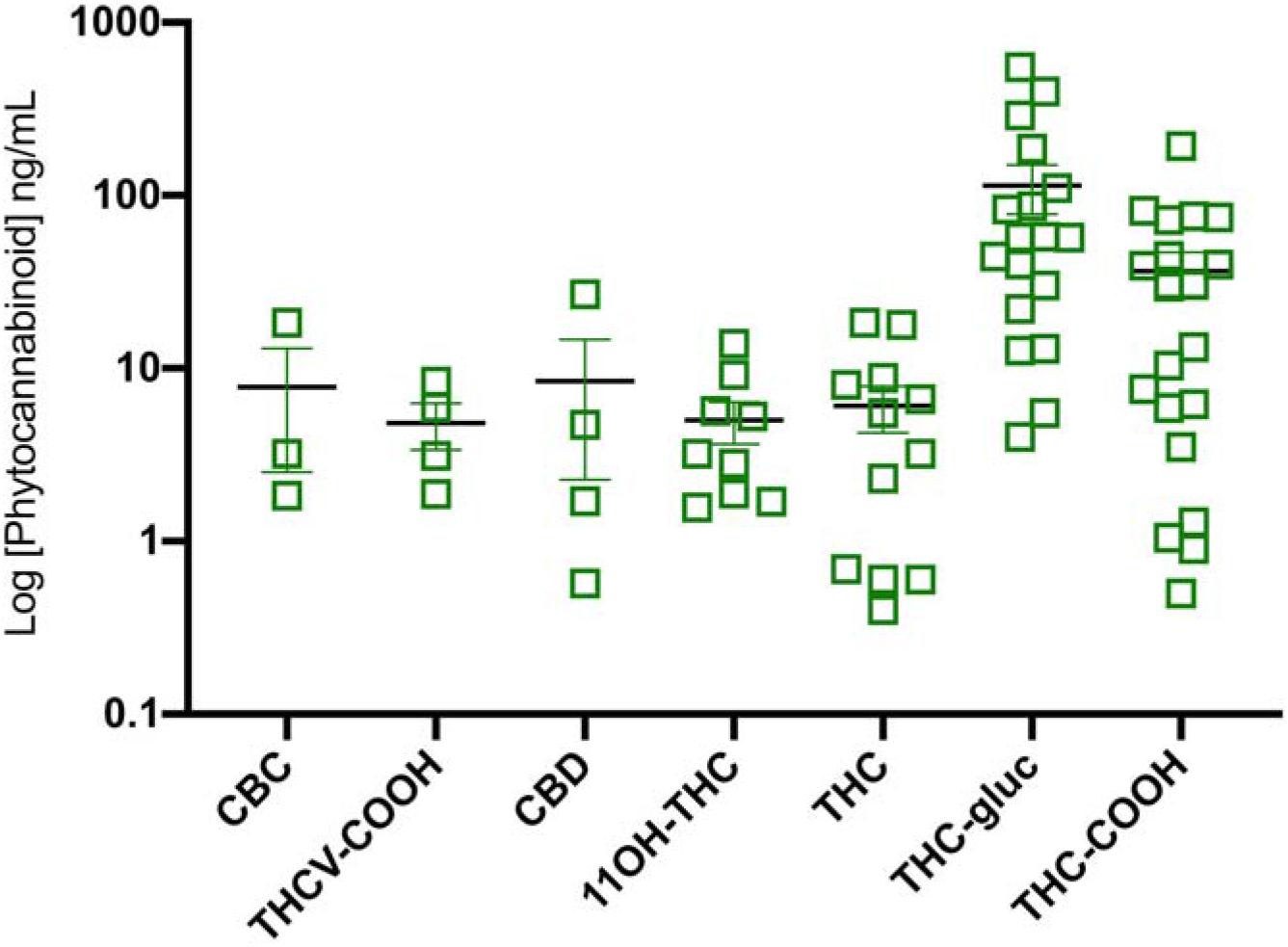
Serum phytocannabinoid profile in adolescent IBD cannabis users. Serum phytocannabinoid levels of adolescent IBD patients enrolled in the CANDID clinical trial who self-identified as having ever used cannabis. Levels are organized by number of patient positive tests starting at cannabichromine (CBC; n = 3) and increasing to THC-COOH (n = 20). Serum levels of cannabinol (detectable range 1.56 to 400 ng/mL), cannabigerol (0.78 to 400 ng/mL), tetrahydrocannabivarin (0.78 to 400 ng/mL) and cannabidivarin (0.78 to 400 ng/mL) not shown here, were all below the limit of detection in all cases. Results represent the mean ± SEM for n= 3-20 patients.

**Figure S2.**
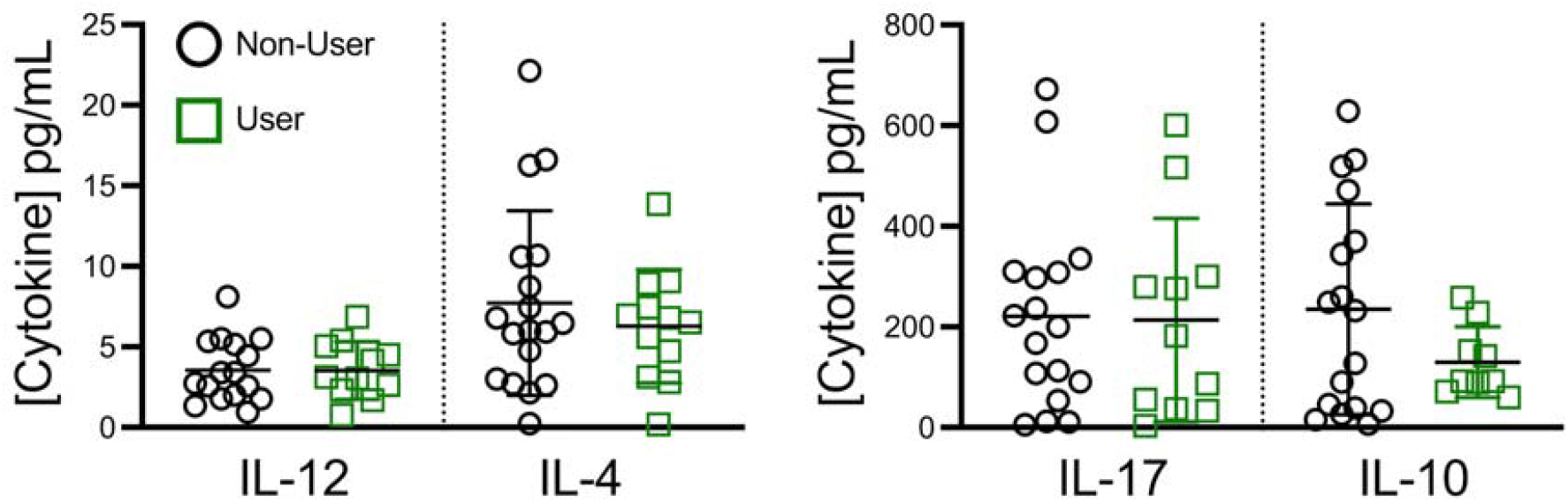
Production of cytokines unchanged in stimulated PBMCs from pediatric IBD patients using cannabis compared to non-users. There was no difference in Th1 (IL-12), Th2 (IL-4), (B) Th17 (IL-17) or Treg (IL-10) associated cytokines between IBD patients who had detectable blood cannabis metabolite levels (“users”) and non-users after 24h restimulation with anti-CD3/CD28 antibodies. Results represent mean±SEM for N ≥ 12.

